# The Structures of Natively Assembled Clathrin Coated Vesicles

**DOI:** 10.1101/2020.01.28.923128

**Authors:** Mohammadreza Paraan, Joshua Mendez, Savanna Sharum, Danielle Kurtin, Huan He, Scott M. Stagg

## Abstract

Vesicle trafficking by clathrin coated vesicles (CCVs) is one of the major mechanisms by which proteins and nutrients are absorbed by the cell and transported between organelles. The individual proteins comprising the coated vesicles include clathrin heavy chain, clathrin light chain, and a variety of adaptor protein complexes. Much is known about the structures of the individual CCV components, but data are lacking about the structures of the fully assembled complexes together with membrane and in complex with cargo. Here we determined the structures of natively assembled CCVs in a variety of geometries. We show that the adaptor β2-appendages crosslink adjacent CHC β-propellers and that the appendage densities reside almost exclusively in CCV hexagonal faces. We resolve how AP2 and other associated factors in hexagonal faces form an assembly hub with an extensive web of interactions between neighboring β-propellers and propose a structural model that explains how adaptor binding can direct the formation of pentagonal and hexagonal faces.

## Introduction

Clathrin-mediated vesicle trafficking is a complex machinery in eukaryotes that is involved in most endocytic pathways (*1*), trafficking from trans-Golgi network (*2*), trafficking from endosomes(*3*), and also in the critical function of regenerating synaptic vesicles (*4-6*). The two main functions of this machinery are engaging with cargo and membrane and also generating negative curvature on membrane such that the cargo ends up in a vesicle. Both of these functions have been shown to be facilitated by the action of adaptor/accessory proteins which are often multi-functional and share overlapping functions (*7-9*). Though clathrin binding has been shown to be dispensable for curvature generation, it plays a major role in organizing adaptors and cargo into spherical vesicles (*10-12*) by making geometrical cages of different sizes both *in vitro* and *in vivo (13, 14)*. Clathrin together with the adaptors AP1 and AP2 act as assembly hubs due to their multiple binding sites for other adaptor/accessory proteins (*15-17*). The dynamic assembly of clathrin-coated vesicles has been investigated by super-resolution time-resolved fluorescence microscopy (*8, 11, 18, 19*), and the self-assembly properties of clathrin have been understood by cryo-EM (*14, 20, 21*). However, it remains to be explained how clathrin always assemble into a fullerene cage with tight geometrical constraints (*22*), in a mechanism that serves to transport cargo in lipid vesicles irrespective of cage geometry. In other words, what kind of interactions coordinate these two aspects such that they never interfere with each other.

Clathrin is a 190 kDa protein with an N-terminal WD40 β-propeller domain (residues 1-330) followed by three coiled-coil domains in the shape of a leg (Fig. S1), and a C-terminal trimerization domain that assembles a clathrin triskelion (Fig. S1), which is the basic assembly unit of the clathrin cage. Clathrin alone can self-assemble to cages *in vitro* in acidic buffer due to multiple salt bridges. However, in physiological pH clathrin light chain inhibits cage assembly. A mechanism was proposed in which assembly was inhibited when the CHC knee residues (R_1161_KKAR_1165_) interacted with CLC’s N-terminal EED motif, which has been shown to be involved in the pH dependence of cage assembly (*21, 23*). For proper assembly, according to this mechanism, CHC’s knee domain can be bent when the interaction with CLC is absent. CLC’s inhibition on assembly is removed *in vivo* by HIP1 (*24*) which interacts with the light chain N-terminal conserved sequence (20-41) via its DLLRKN-containing coiled-coil domain. Additionally, CLC also has a central heptad repeat that interacts with the CHC proximal domain, and a C-terminal domain that interacts with the tripod helices. It has been shown through FRET measurements that light chain heptad repeat undergoes a conformational change upon cage assembly. Additionally, it has been shown through atomic force microscopy that the light chain rigidifies the CHC cage (*25*).

CHC β-propellers provide the grooves between their blades for binding of most adaptor/accessory proteins (*26*) like AP1-3, AP180, HIP1, and EPSIN which possess multiple linear motifs (clathrin box) in their C-terminal unstructured regions. AP1 and 2 also have appendages (β1 and β2, respectively) C-terminal to their clathrin boxes that have been shown to bind clathrin (*16, 27, 28*). Although clathrin box binding sites are well known, the appendage binding site has been mapped to a broad region of the N-terminal CHC coiled coils (*29*). Since AP2 also binds other adaptor/accessory proteins as well as PIP4 and cargo recognition peptides, these features make AP2 (and its counterpart AP1 in Golgi/endosome trafficking) the nucleator of CCV assembly and the main recruiter of clathrin. Though its binding to clathrin has been well established, it has remained unclear how AP2 can structurally organize the formation of coherent spherical clathrin cages.

Clathrin is shaped such that it assembles into pentagonal, hexagonal, and rarely heptagonal faces. There are always 12 pentagons in a cage (13 in the rare cases of cages with heptagonal faces). As with any fullerene structure, curvature is only generated by the presence of pentagonal faces, and the diameter dictated by the number of hexagonal faces. For the clathrin cage, in any given edge, four different CHCs interact extensively along their proximal and distal domains. A 4.7 Å structure of clathrin cages was published recently that revealed that the interactions among different triskelion legs are highly conserved regardless of the cage size or face geometry (*14*). In that study, structures were determined of *in vitro* assembled cages together with the β2-adaptin of AP2, but notably, they did not observe the β2-adaptin density. All adaptor-clathrin interactions have been mapped to the N-terminal domain of clathrin, a region that is far from the extensively interacting proximal and distal domains. Thus, a remaining question for the field is what is the interplay between adaptor binding and the generation of cages of given diameters? How are the number of pentagonal and hexagonal faces determined? Here using cryo-EM, we have determined the structures of natively assembled CCVs that include clathrin, adaptors, accessory proteins, and cargo. Our structures show a number of previously unresolved features including adaptor β2-appendages that crosslink adjacent CHC β-propellers. Remarkably, the appendage densities reside almost exclusively in hexagonal faces, and by performing a focused alignment and classification, we resolve how AP2 and other associated factors in hexagonal faces form an assembly hub with an extensive web of interactions between neighboring β-propellers. These observations lead us to propose a structural model that explains how adaptor binding can direct the formation of pentagonal and hexagonal faces.

## Results

### Natively assembled CCVs assume 5 different cage geometries including one not thought to be possible

Clathrin-coated vesicles were purified from bovine brains in conditions that ensured CHC cages do not disassemble during purification. SDS-PAGE gels showed bands that corresponded to clathrin heavy chain and light chain and also adaptor subunits (Fig. S2A). The identities of these proteins and other accessory and cargo proteins were determined by mass spectrometry (Fig. S2B). This showed that the adaptors AP2 and AP180 were highly abundant followed by AP1 and AP3. Other essential accessory proteins were present in the CCVs including AP2-related kinase, EPSIN, HIP1, and amphyphisin. Besides AP2 and AP180, all the essential components of synaptic vesicles (SVs) (*30*) were present in the spectra, including: VAMP2 (R-SNARE protein), synaptophysin (VAMP2 chaperone and inhibitor), synapsin (localizes SVs to the synapse pool), sodium-potassium ATPase pump and H^+^ ATPase pump, and neurotransmitter transporters (Fig. S2A). SV-related proteins had the highest number of unique peptides and total normalized spectra in the mass spectrometry analysis reflecting their higher concentration in the purified CCVs. Among other localized proteins are syntaxin 1 and 2 (plasma membrane), syntaxin 7 (early endosome, late endosome), and syntaxin 16 (*trans*-Golgi network).

The natively assembled CCVs were prepared for cryo-EM and a single particle cryo-EM dataset was collected resulting in 80k CCV particles (Fig. S2B). These were subjected to RELION’s ab initio model generation (*31*) with a target of 4 models. This resulted in four CCV structures with complete cages and unique symmetries including the mini-coat (tetrahedral, 28 triskelions), football (D3, 32 triskelions, also known as sweet potato (*14*)), tennis ball (D2, 36 triskelions), and D6 barrel (D6, 36 triskelions), (Fig. 1A). However, by inspection of the individual particles it was clear that there were more geometries present in the dataset. Therefore, the cages were subjected to multiple rounds of competitive 3D classification where the particles that were already assigned to one class were further classified. This led to the identification of a subset of particles that form a previously undescribed geometry with C2 symmetry and 30 triskelions that we are calling the C2 basket. The mini-coat, football, tennis ball, and barrel structures match four of the cage geometries that were recently reported by Morris *et al*. (*14*) from *in vitro* reconstituted clathrin cages where they used a library of pre-generated cage geometries to sort their particles into the representative cage structures. This indicates that the cellular processes that assemble CCVs are similar to clathrin cages generated by reconstitution. However, there are a few exceptions; with our data-driven model generation approach, we were not able to unambiguously generate the C1 cage that Morris *et al*. observed by refining against pre-generated cage geometries. It is possible that the C1 cages are in our dataset, but they are likely such low abundance that we cannot unambiguously identify their presence. Another difference is that our largest identified cage (36 triskelions) is the smallest among 8 reported cages by Cheng *et al*. (*13*) that were purified as intact CCVs similar to our approach. Although intact CCV sample preparations follow the same ultracentrifugation protocols with slight modifications, the lack of cages larger than 36 triskelions in our sample and the lack of cages smaller than 36 triskelions in their sample indicates that either the source tissues differ in some way or that a subtlety in methods leads to the enrichment of smaller or larger CCVs. We have high confidence that our CCVs are representative of synaptic vesicles due to the abundance of SV proteins identified by mass spectrometry.

**Figure 1.**
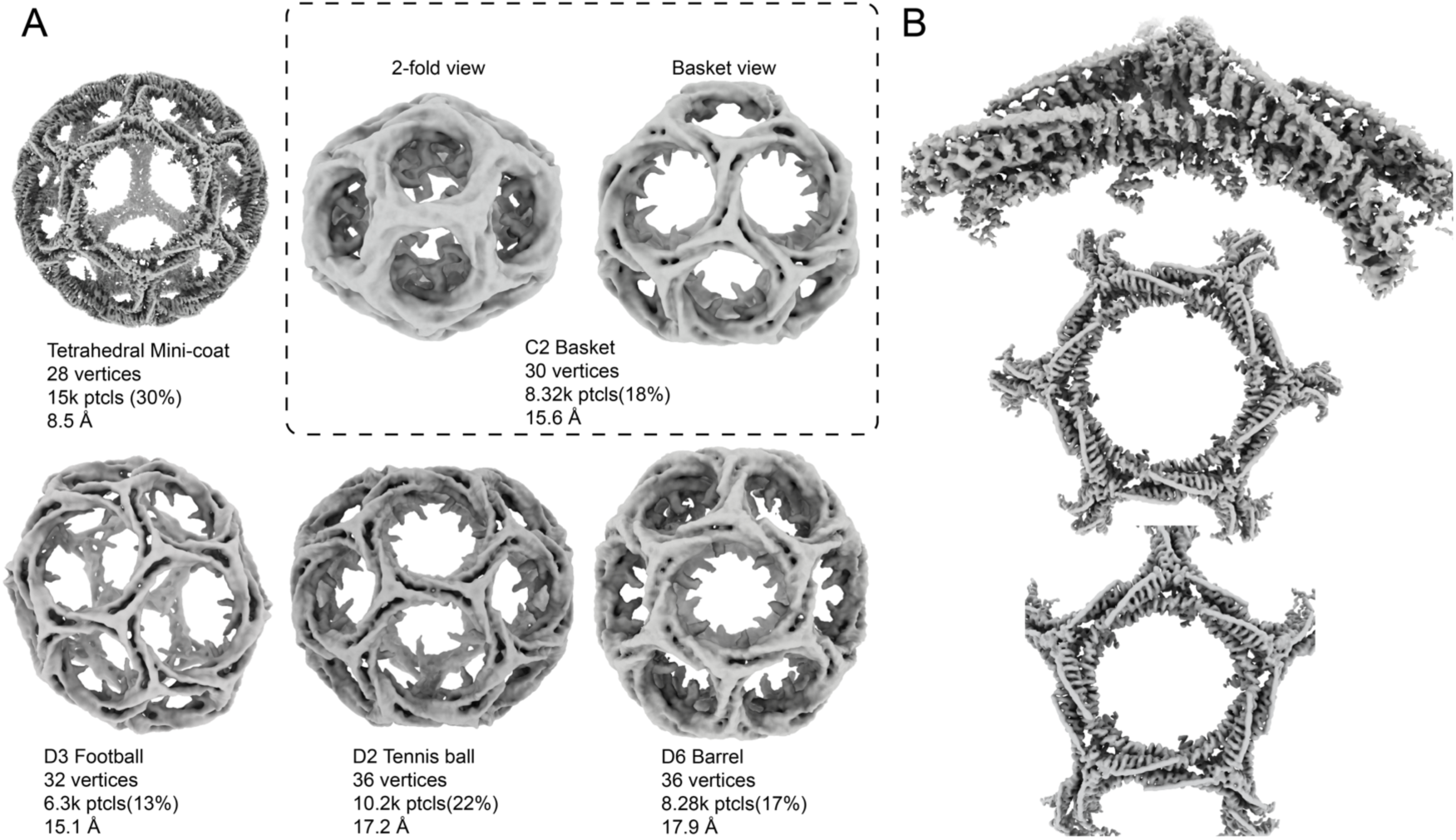
Refinement of the clathrin layer of CCVs. A) Refinement of whole clathrin cages into 5 different classes. The tetrahedral mini-coat has the highest resolution among cages. The C2 basket is a novel geometry represented in two views in the dashed panel. The largest cage is the D6 barrel. B) Clathrin-focused subparticle refinement of the mini-coat structure further improves the resolution. Top: asymmetric vertex refinement, resolution: 6.3 Å. Middle: hexagonal face refinement, resolution: 9.1 Å. Bottom: pentagonal face refinement, resolution: 8.27 Å.

The C2 cage that we observe by cryo-EM is novel in two ways: first, it has not been reconstructed before in cryo-EM datasets, second, its existence was ruled out by a cage geometry prediction model developed by Schein (*22*) and utilized by Morris *et al*. (*14*) in structure determination. This model uses a so-called head-to-tail exclusion rule to find possible cage geometries given the number of vertices. This rule stems from avoiding structures with too much dihedral angle discrepancy that would cause the polygonal faces to be non-planar. According to this model, fullerene cages with 30 vertices are not permitted at all. In contrast, our C2 structure consists of 30 vertices, 12 pentagons, 5 hexagons, and 45 edges. Although this structure violates the head-to-tail exclusion rule, it follows Euler’s polyhedron formula which leads to the formation of closed polyhedrons. The specific region of this geometry that violates the head-to-tail rule consists of 4 adjacent pentagons with 5 mutually-shared edges (Fig. S3A). Two of these pentagons each contain two vertices where the rule is violated and thus there should be excessive torsion (Fig. S3A). This explains the slightly weaker density for the CHC chains highlighted in Figure. S3A.

The refined clathrin cages were further subjected to subparticle refinement (*32*) to improve the resolution of the clathrin layer. The asymmetric vertex (surrounded by hexagonal-pentagonal-pentagonal or HPP faces) of the mini-coat refined to 6.3 Å (FSC_0.143_) overall with some areas as high as 4.6 Å (Fig. 1B, Fig. S4D), while its pentagonal and hexagonal faces refined to 9.1 and 8.3 Å, respectively (Fig. 1B). The other cage geometries also produced subparticle reconstructions for the HPP asymmetric vertex with resolution ranging between 8-9 Å (Fig. S4). This held true for the asymmetric vertex of the C2 basket structure as well (Fig. S3, Fig. S4).

### Atomic modeling of the asymmetric vertex structure reveals interactions of the QLMLT motif of the CHC and the stabilizing interactions of the CLC with CHC

Atomic models were built of the mini-coat HPP asymmetric vertex. The model by Morris *et al*. (*14*) that was built into their consensus C3 vertex was docked into our C1 vertex (Fig. 2B). In this rigid-body fit, the edge of the vertex that is between two pentagonal faces exhibits an accurate fit, while the two other edges with one hexagonal face do not enclose the model starting at positions close to the knee domain (Fig. 2B). This is consistent with observations that the CLC stabilizes the proximal domain and that the knee domain changes conformation to make hexagonal and pentagonal faces (*14*). Using the asymmetric vertex structure from the mini-coat (*14*), a homology model of the bovine CHC and CLC was built and refined to our map (Fig. 2A). 15 C-terminal residues of CHC were built *de novo* including QLML residues of the QLMLT motif (*33*). These residues follow the final C-terminal helix of CHC at the trimerization domain which itself sits on top of three distal legs crossing below making a triangle (Fig. 2C). We show that the QLML motifs then extend from the trimerization helices such that each motif makes interactions with two crossing distal legs (Fig. 2C). In this way, three motifs stabilize three pair-wise interactions between three CHC chains. Another important interaction that happens at the center of the vertex is between CHC and CLC. Three C-terminal helices of CLC form a U-shaped domain that lies against CHC residues at the trimerization domain burying hydrophobic residues along the interface and stabilizing the tripod domain (Fig. 2D).

**Figure 2.**
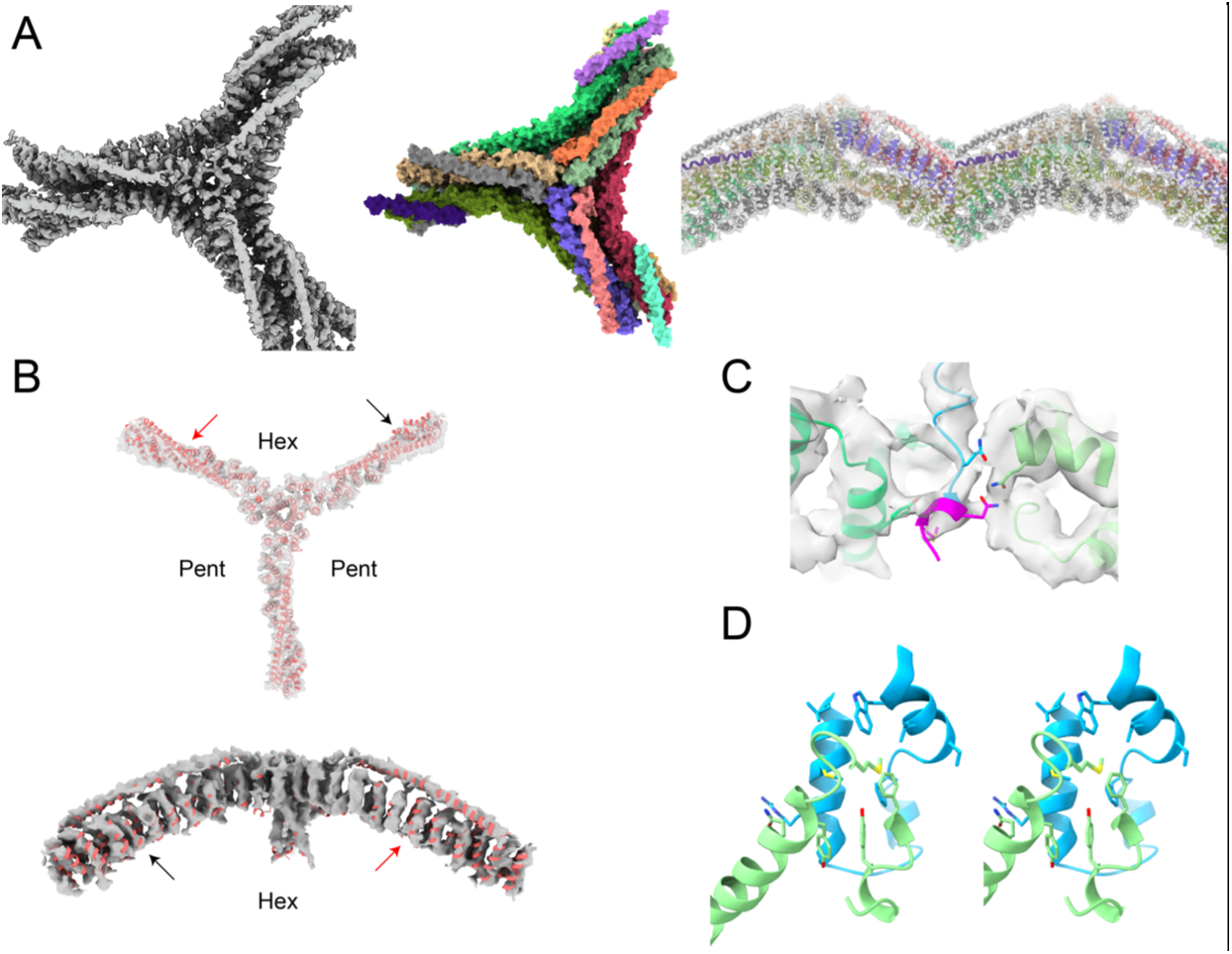
Atomic modeling of the asymmetric vertex of the mini-coat. (A) The left panel shows the vertex structure used for building the atomic model. The middle panel shows the atomic model colored according to different CHC and CLC chains. The right panel is a stereo view of the model fitted in the map in a view perpendicular to that of the left and the middle panels. B) PDB-6SCT by Morris *et al*. fitted in our asymmetric vertex map. Top panel: red arrow shows the point in the clathrin chain that curves inside the hexagonal face where the model does not fit the map anymore, black arrow shows the deviation point for the clathrin chain that curves inside a pentagonal face. Bottom panel: the same deviation points are indicated with arrows for a view that is perpendicular to that of the top panel. C) one QLML sequence shown in magenta from the trimerization helix interacts with two distal legs shown in light green. D) stereo view of the C-terminal domain of CLC shown in cyan lying against CHC residues shown in light green.

### Adaptor appendage densities attached to CHC β-propellers are enriched in hexagonal faces

Inspection of the CHC β-propeller densities in the mini-coat reconstruction revealed that there is a density consistent with the AP2 appendage domains that resides exclusively in the hexagonal faces and not in the pentagonal faces (Fig. 3A). Docking the crystal structure of the β2-appendage (*34*) in the observed densities aligns residues Y815 and Y888 of the two appendage domains (*16, 35*) with two neighboring β-propeller domains from two different CHCs (Fig. 3A) effectively crosslinking the two CHCs. For each β2-appendage, one of the two domains connects down to the body of the adaptor which allowed us to orient the domains. In the tetrahedral mini-coat, the hexagonal faces lie on three-fold axes of symmetry, and three appendages can easily be accommodated by the six β-propellers in the hexagonal face. The other cage geometries also have an abundance of appendages in hexagonal faces with the hexagonal faces showing two or more appendage densities (Fig. S5). With the larger geometries, weak appendage densities could also be observed in pentagonal faces that have multiple neighboring hexagonal faces as discussed in the following sections.

**Figure 3.**
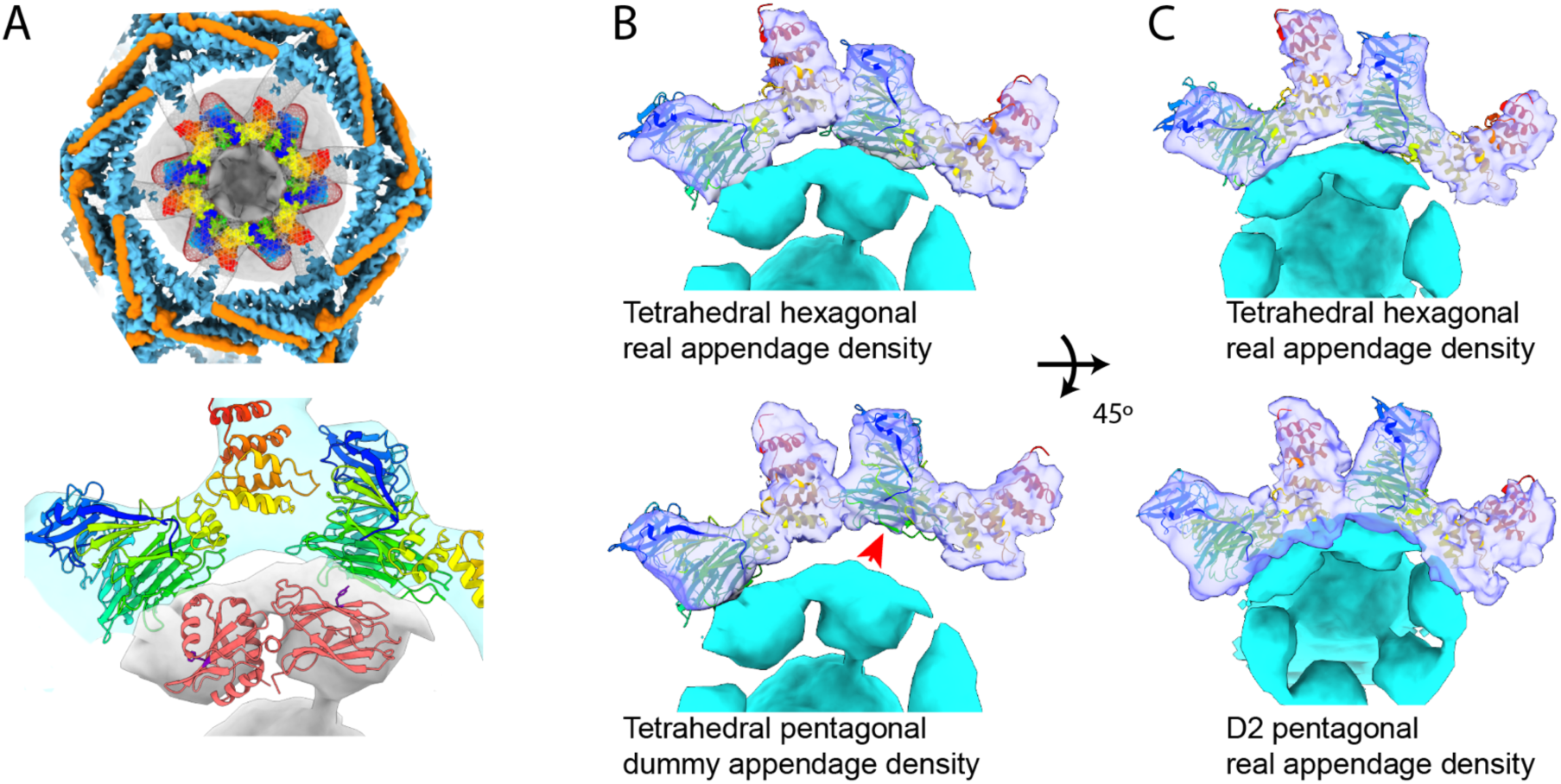
Three-way interaction between β-propeller and β2-appendage and β-propeller and ankle. A) CHC and CLC are indicated in light blue and orange, the β-propeller and part of ankle crystal structure (PDB-1BPO) colored red to dark blue corresponding to C-terminus and N-terminus, and β2-appendage density shown in grey. Top panel: ankle and β-propeller densities are shown in mesh with crystal structure of β-propeller and part of ankle fitted inside. Three β2-appendage densities are resolved in the hexagonal face. Bottom panel: zoomed in view of β2-appendage (crystal structure PDB-1E42 shown in pink) interacting with two β-propellers with the side chains of two crucial tyrosine residues represented. B,C) Ankle and β-propeller map are represented in transparent purple, appendage density is shown in cyan. B) top panel: relative orientation of two neighboring β-propeller and ankle domains in the mini-coat hexagonal face, the left leg and the appendage density remain in place between top and bottom panel. Bottom panel: in the pentagonal face the right leg slides on the neighboring ankle away from the dummy appendage density (indicated with red arrow). C) top panel: top view of top panel in B. Bottom: top view of bottom panel in B, however, the appendage density is real (D2 cage) and it has slid under the β-propellers and changed orientation.

Compared to the reconstituted clathrin cage reconstructions by cryo-EM (*14, 20, 36*), our structures have strong densities for the CHC β-propeller domains, and they are always in contact with the neighboring CHC ankle domain. We observe two preferred conformations for these contacts that differ significantly between the hexagonal and pentagonal faces (Fig. 3B) and are consistent for all cage geometries. In the hexagonal faces, the β-propeller interacts with the N-terminal end of the ankle near residues 330-480. In pentagonal faces that have adaptor appendage densities, the appendages bind the neighboring β-propellers the same as in the hexagonal faces, but because pentagonal faces are smaller the β-propellers shift up the ankle and below the β-propellers in order to preserve the appendage-β-propeller interaction (Fig. 3C).

### Pentagonal faces with adaptor appendages have at least two neighboring and consecutive hexagonal faces

Unlike hexagonal faces where adaptor appendages are found in abundance in all geometries, appendage densities in pentagonal faces depends on their neighboring faces. In the five resolved cage geometries, there are three possible configurations of neighboring faces for each pentagonal face (Fig. 4A-C). The first configuration has two non-neighboring hexagonal faces and appears in the T (mini-coat), D3 (football), and C2 (basket) geometries (Fig. 4A, top panel). This configuration has no associated adaptor appendages in the pentagonal faces (Fig. 4A, bottom panel). The second configuration is unique to C2 geometry and has two neighboring hexagonal faces that adjoin each other (Fig. 4B, top panel). In this configuration two of the β-propellers in the pentagonal face are interacting with adaptor appendages (Fig. 4B, bottom panel). The third configuration has three total hexagonal faces two of which adjoin each other and appears in D6 (barrel) and D2 (tennis ball), D3, and C2 geometries (Fig. 4C, top panel). In this configuration all β-propellers in the pentagonal faces interact with adaptor appendages (Fig. 4C, bottom panel). This observation clearly shows that as the number of hexagonal faces and hence the cage size increases, more pentagonal *β*-propellers engage with adaptor appendages. Moreover, since the number of pentagonal adaptor appendages correlates with the number of neighboring hexagonal faces, this suggests that a cluster of adaptors below hexagonal faces also engage their neighboring pentagonal β-propellers. This idea was borne out by performing, subparticle analysis of hexagonal faces and pentagonal faces’.

**Figure 4.**
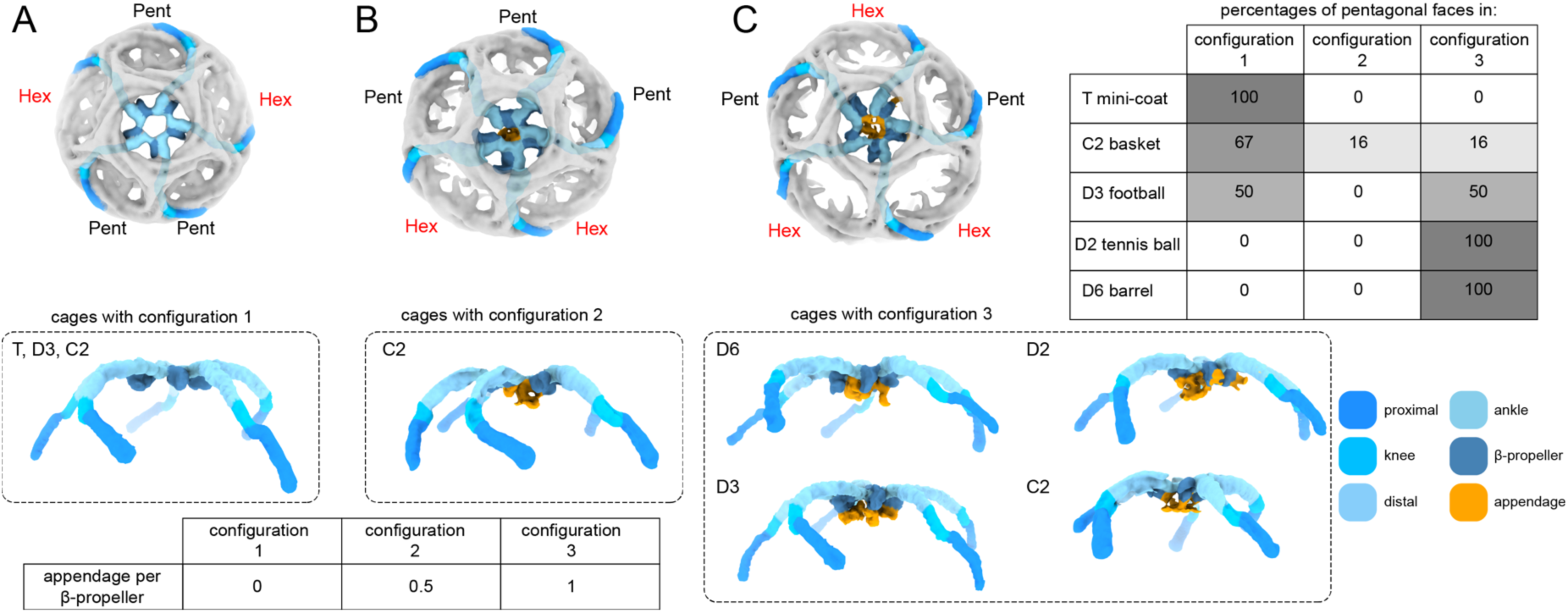
Three different configurations of neighboring faces for a pentagonal face in all geometries and their relation to the population of adaptor appendages in the pentagonal face. The color code at the bottom right corner applies to all. A) Top panel: Configuration 1 for faces neighboring a pentagonal face. The neighboring faces are indicated with Pent and Hex. The CHC chains with β-propellers in the central pentagonal face are traced using the color code. Bottom panel: The same CHCs traced in the top panel is shown in a side view with no appendages present. Geometries with configuration 1 are indicated (T, D3, C2).B) Top and bottom panels of configuration 2 are arranged in the same manner as in A. This configuration is unique to C2 basket. Bottom panel shows that some β-propellers are engaged with adaptor appendages. C) Top and bottom panel of configuration 3 are arranged in the same manner as in A. Four different geometries indicated in the bottom panel accommodate configuration 3 in which all β-propellers are engaged with adaptor appendages. The top table shows the abundance of the three configurations in 5 different geometries. The bottom table shows relationship between configuration and approximate percentage of β-propellers in the central pentagonal face of that configuration that are engaged with adaptor appendages.

### Subparticle refinement reveals AP2 density

Since adaptor densities were present in our CCVs but we were only resolving the appendages, we performed subparticle refinement on the hexagonal faces to better resolve the adaptors. 60k hexagonal face subparticles were generated from 15k mini-coat CCVs. A focused classification scheme was used that enabled alignment of the density below CHC to a greater extent (Fig. 5A). This revealed 8 hierarchical classes that all showed adaptor density, presumably AP2, in the center of the hexagonal faces. Although the AP2 density was relatively low-resolution (Fig. 5B), it is distinguished by 3 features (Fig. 5C): 1) the core AP2 complex in the open conformation (PDB-2XA7, (*7*)) with the hetero-tetramer of α, β2, σ2, and μ2 subunits fitting inside the corresponding AP2 density, 2) AP2 density makes two points of contact with membrane corresponding to the reported α and μ interactions with PIP4 and cargo peptides, respectively, 3) appendage densities connect AP2 to CHC β-propeller domains (Fig. 5D). Additionally, there are multiple densities extending from the AP2 density that engage the β-propellers of neighboring pentagonal faces (Fig. 5D,E). These observations are consistent with the idea that hexagonal adaptor clusters can engage more than just six hexagonal β-propellers. Our hexagonal face map shows that a total of 18 β-propellers are engaged with a cluster of proteins centered on the AP2 adaptor (Fig. 5E).

**Figure 5.**
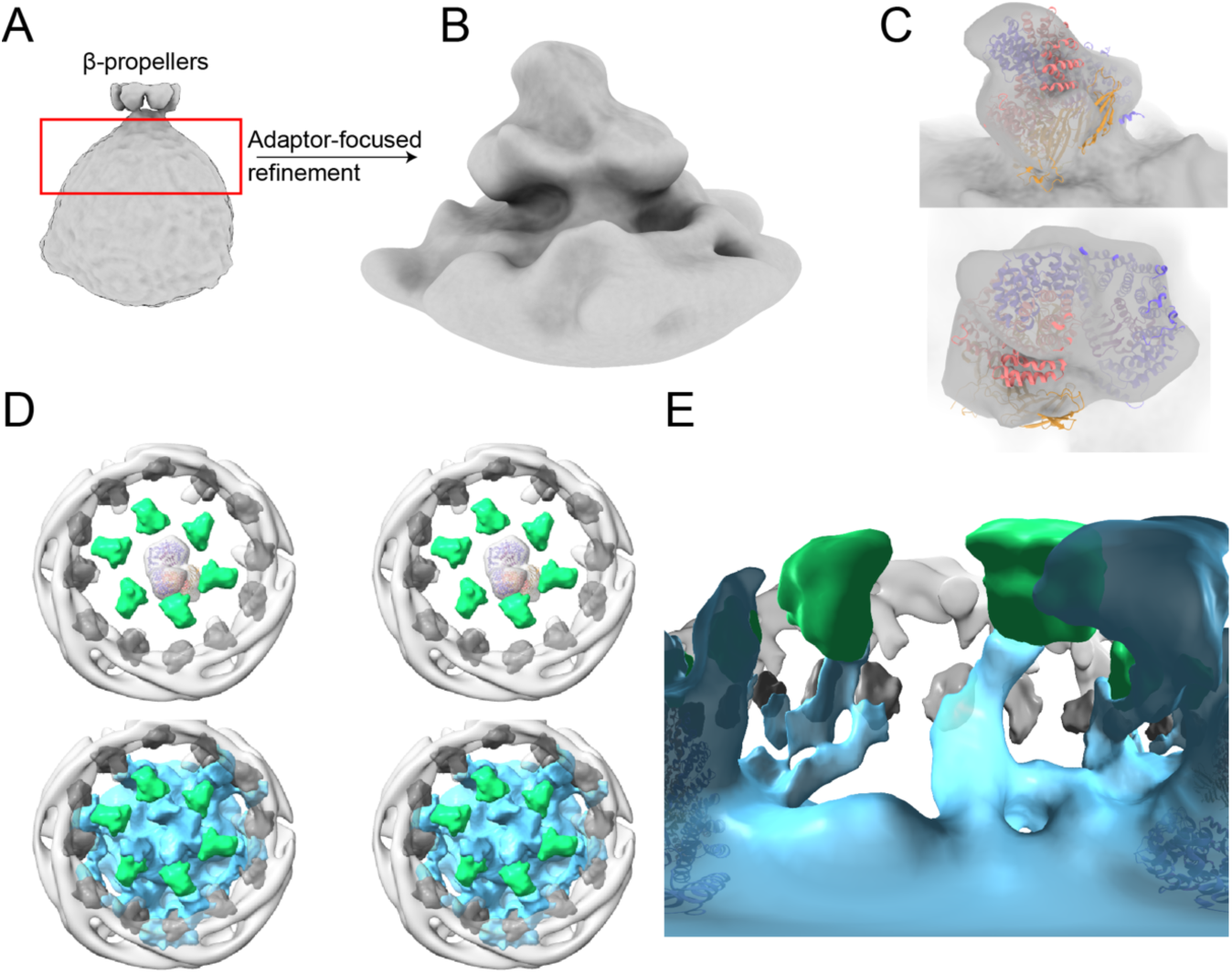
Resolving AP2 density of the mini-coat hexagonal face and a crowded web of appendages. A) β-propellers of one hexagonal face and averaged-out density from the mini-coat cage refinement, red box indicates the volume that was further resolved in subparticle alignment. B) the result of subparticle alignment reveals a density resembling the AP2 core. C) crystal structure of open AP2 fitted inside the AP2 map. D) β-propellers of the hexagonal face (green) and the neighboring pentagonal faces (dark grey), total of 18, are represented along with some of the CHC proximal domain densities to show orientation. Top panel (stereo view): AP2 core density is indicated in transparent pink with the crystal structure fitted inside. Bottom panel (stereo view): at a different density threshold 17 β-propellers become engaged with what seems to be different adaptor appendages and adaptor C-terminal domains. E) 360° view with the adaptor eye looking at hexagonal and pentagonal β-propellers and their interactions with adaptors. Colors and crystal structure are the same as in D.

## Discussion

Structure determination of natively assembled CCVs has revealed many new insights into the interplay between adaptor binding and clathrin cage assembly. CCVs are highly heterogeneous in size, shape, and composition especially with respect to the membrane and cargo proteins (Fig. S2B). Nonetheless, using cryo-EM we were able to classify the cages into five discrete geometries that represent the majority of CCVs we imaged. Using subparticle reconstruction, we determined the structure of the most abundant vertex in our dataset, which is asymmetric and formed form the intersection of two pentagonal faces and one hexagonal face. This structure confirmed that the interactions that Morris *et al. (14)* observed in their reconstituted clathrin cages were conserved in our natively assembled coats. Similarly to their study, our structure also showed that the CLC C-terminal domain stabilizes the tripod domains that are critical for CHC trimerization. A unique feature of our structure was the C-terminal residues of the CHC tripods that extend from the base of the tripod helix past the distal legs of the three CHCs beneath the vertices such that the CHC legs are linked together through the tripod helices (Fig. 2C). The tripod helices extend a bit past this region partially exposing the QLMLT motif that is recognized by Hsc70 (*33*). These observations explain how Hsc70 can compete with the tripod/distal domain interaction to destabilize cages.

One of the most striking features of the structures we observed was the enrichment of adaptor densities in the hexagonal faces of the CCV structures. In our most abundant cage, the mini-coat, there were strong densities for AP2 β2-appendages exclusively in the hexagonal faces with no noticeable density in the pentagonal faces. As the cages increased in diameter, adaptor density increased in the pentagonal faces, but only for faces that were bounded by two adjoining hexagonal faces (Fig. 4). We discovered that individual appendage domains crosslink the two β-propellers of two neighboring CHCs, which in turn stabilizes a previously uncharacterized interaction between the β-propeller of one CHC and the ankle of its neighboring CHC (Fig. 3). In this way the binding of a single AP2 ties together two triskelions and promotes cage assembly. This observation also explains why reconstructions of *in vitro* reconstituted cages lacking β2-appendages do not show the same ubiquitous β-propeller-ankle interaction that we observe. When we performed an asymmetric subparticle focused refinement on the hexagonal faces, we resolved AP2 and a whole network of interacting densities that link the hexagonal face to the 12 CHC β-propeller domains in neighboring faces (Fig. 5). In this way AP2 and other associated factors can serve as a hub linking cargo binding, adaptor assembly, and clathrin assembly.

All of these observations have led us to propose a model for how adaptor binding drives the formation of discrete cage geometries. It is well-established that adaptors like AP2 drive curvature on membranes. Our data demonstrate that β2-appendages are enriched in hexagonal faces and crosslink neighboring clathrin triskelia thereby promoting the assembly of hexagonal faces. However, for the formation of a closed cage there should be a mechanism that explains how pentagonal faces are formed in the two modes of assembly: constant area and constant curvature (*11, 37*). In the constant curvature model, pentagonal faces would form in membrane regions lacking adaptor clusters in order to accommodate membrane curvature. For the constant area model, CCVs start out as a flat hexagonal lattice, and triskelia have to be removed by Hsc70/auxillin to convert hexagonal faces t o pentagonal ones (*38-40*). Our data suggest that the presence of an adaptor cluster in a hexagonal face prevents the removal of that face’s triskelia through multiple appendage-β-propeller interactions (Fig. 6, gray). In this view, these triskelia would remain even if the tripod interactions were disturbed temporarily by Hsc70/auxillin. Furthermore, in the neighboring hexagonal faces, there is only one triskelion that is not engaged in any adaptor hub interactions (Fig. 6, red). If all of those triskelia were removed by Hsc70/auxillin (Fig. 6, green), then the adaptor cluster would give rise to a hexagon surrounded by six pentagons thus accommodating a curved membrane. In order to make larger cages, more hexagonal faces must be incorporated via adaptor clusters with the same pattern of appendage-β-propeller interactions (Fig. 6). This is consistent with our observation that the D2 tennis ball and D6 barrel that have multiple consecutive hexagonal faces also have pentagonal faces that are enriched in appendage densities (Fig. 4). Thus, in our model, the positioning of the 12 pentagons that are required for the generation of a closed geometrical lattice are not determined by the geometrical constraints of clathrin but rather the positioning of adaptors. This central role of adaptors in generating clathrin geometries is supported by our observation of the C2 basket, which should be unlikely to form based on clathrin interactions alone. In this way, our model unifies the literature on CCV assembly by describing clusters of adaptors as the central machinery for curvature generation and CHC cage assembly.

**Figure 6.**
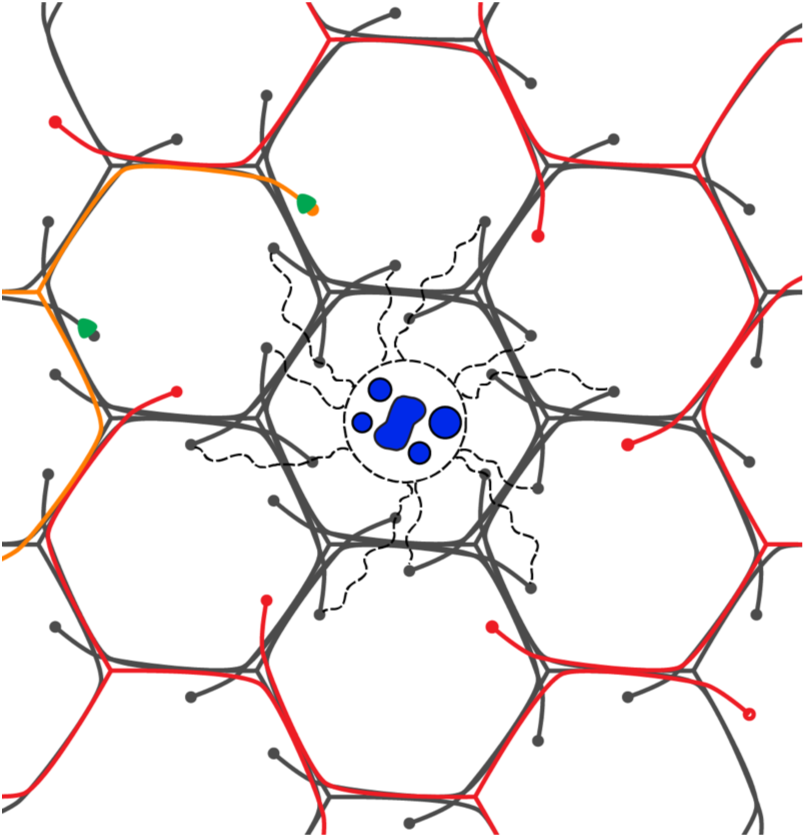
Model for adaptor mediated formation of discrete clathrin geometries. An adaptor cluster in the center of a hexagonal clathrin face (blue) links the hexagonal face to the 12 CHC β-propeller domains in neighboring faces stabilizing their triskelions. The triskelions in red represent ones are candidates for removal in order to generate a pentagonal face. The orange triskelion is actively being removed by Hsc70/auxillin (green)

## Accession codes

All cryo-EM maps and structures are being deposited in the Electron Microscopy Data Bank and Protein Data bank with IDs xxx.

## Acknowledgements

The authors acknowledge the use of instruments at the Biological Science Imaging Resource supported by Florida State University and NIH grants S10 RR025080 and S10 OD018142. The work presented was supported by the NIH grant R01GM108753 and funds from Florida State University.

## Methods

### Purification

All the following steps were done at 4°C. 120g of cow brain purchased from Pel-Freez Biologicals was homogenized in MES buffer (100mM MES, 1 mM EGTA, 0.5 mM MgCl2, pH 6.7) (*41*) plus 2 protease inhibitor cocktail tablets using an industrial blender (Final volume 300 ml). The homogenate was centrifuged at 17k *g* and the supernatant was transferred and centrifuged at 56k *g*. The pellet from that run was resuspended in MES buffer plus %12.5 w/v of each sucrose and ficoll and then homogenized again using the blender. The homogenate was spun at 43k *g* to pellet large microsomes. The supernatant was transferred and spun at 100k *g*. The pellet was resuspended in MES buffer and spun at 17k *g* to remove any remaining unwanted large lipids. The intact clathrin coated vesicles (two tubes of 7.5 ml) were then spun on an %8 sucrose cushion dissolved in MES buffer made with D_2_O (2.5 ml, added to the bottom of the sample) at 116k *g* to separate out similar-sized uncoated vesicles. The pellet was resuspended in MES buffer at a concentration of 6.5 mg/ml.

### Mass spectrometry

#### Protein digestion

Cut gel bands were de-stained with a wash buffer (50% aqueous acetonitrile with 50 mM ammonium bicarbonate) and cut into ∼ 1 mm gel pieces and dried by incubation with acetonitrile followed by SpeedVac (Thermo Fisher Scientific). Digestion buffer (10% aqueous acetonitrile with 50 mM ammonium bicarbonate) was added to rehydrate the dried gel pieces, which subsequently were reduced with 0.5 mM TCEP (tris(2-carboxyethyl)phosphine, Sigma-Aldrich, catalog #: C4706, St. Louis, MO) and alkylated with 1 mM IAA (iodoacetamide, Sigma-Aldrich, catalog #: I1149, St. Louis, MO). Then trypin (Thermo Fisher Scientific, catalog #: 90058, Waltham, MA) was added to a final concentration of 0.005 µg/µL, and the mixture was incubated at 37°C overnight. Digestion was quenched by adding 0.5% formic acid aqueous solution and the supernatant was collected. The residual gel pieces were dried by incubation with acetonitrile and the supernatant was collected. Combined supernatant containing tryptic peptides was dried in a SpeedVac.

For the in-solution protein trypsin digestion, 24.5 µL of the digestion buffer (10% aqueous acetonitrile with 50 mM ammonium bicarbonate) was added to 0.5 µL of the crude CCV sample. The mixture was then reduced with 0.5 mM TCEP, alkylated with 1 mM IAA and digested with 0.005 µg/µL trypsin at 37°C for 3 hr. The digestion was quenched by adding 0.5% formic acid aqueous solution and the tryptic peptide mixture was dried in a SpeedVac.

#### Tryptic peptides separation and detection

The peptide mixture was re-dissolved in 0.1% aqueous formic acid and separated by nano-liquid chromatography (nLC) with an Easy Nano LC II system (Thermo Fisher Scientific) with a 100 µm x 2 cm trap column (easy column, catalog # SC001 Thermo Fisher Scientific) and a 75 µm x 10 cm C18AQ analytical column (Thermo Fisher Scientific). Mobile phases compositions are A (0.1% aqueous formic acid) and B (99.9% acetonitrile and 0.1% formic acid). For the crude CCV sample, a linear gradient from 5% to 35% B over 2 hr was performed with a flow rate of 300 nL/min. For gel band samples, the linear gradient was from 5% to 35% B over 1 hr. Eluate was on-line ionized by nano-electrospray ionization (nESI) with a spray voltage of 2.2 kV and detected by a Hybrid Velos LTQ-Orbitrap Mass Spectrometer (MS) (Thermo Fisher Scientific). Both nLC and MS were controlled by the Xcalibur software (Thermo Fisher Scientific). The precursor ions were detected with a mass resolution of 60 K (at *m/z* of 800 Da) and Automatic Gain Control (AGC) of 1.00e+6 in the Orbitrap. Centroided data-dependent MS^2^ was carried out on the top 10 most abundant precursor ions with collisional induced dissociation (CID) in the LTQ with AGC of 1.00e+4.

#### Data Analysis

Acquired Xcalibur.raw files were analyzed by Proteome Discoverer software (version 1.4) with a Sequest HT search engine (Thermo Fisher Scientific) against a *Bos taurus* protein fasta database with dynamic modification of methionine oxidation and static modification of cysteine carbamidomethylation. The search result was further validated and visualized with Scaffold software (version 4.3.4, Proteome Software Inc., Portland, OR).

### Cryo-EM data collection and preliminary processing

3 μl of sample was applied to thick C-flat grids (CF-2/2-4C-T-5), blotted (blotting time 4s, blotting force 2) and plunged in liquid ethane using a Mark IV vitrobot (chamber humidity 100% at 4°C). Movie frames were collected on a Titan Krios equipped with a K3 direct electron detector, at 33k magnification, super-resolution mode with a pixel size of 1.37Å, a total dose of 40.98 e^-^/Å^2^, and a defocus range of 3-5 microns. 800 micrographs were obtained after motion correction and dose weighting using MotionCorr2 (*42*). Contrast transfer function was estimated using CTFFIND v4 (*43*). CCV particles were picked using FindEM Template picking (*44*), resulting in a total of 80k CCV particle images (894-pixel box size). FSC was generated using even and odd masked half maps, and the local resolution for the mini-coat vertex was calculated using monores (*45*).

### Cage Structure determination

The pool of 80k CCV particle images was 2D-classified using CRYOSPARC (*46*) leaving 60k particles which were then 3D-classified using RELION (*31*) and CISTEM (*47*) into 5 different clathrin cage geometries:

1. *mini-coat* (15k particles, T symmetry, 28 triskelions, 24 C1 (HPP) vertices, 4 C3 vertices, 4 hexagonal faces)
2. *C2 basket* (8.32k particles, C2 symmetry, 30 triskelions, 22 C1 HPP vertices, 4 C3 vertices, 4 C1 vertices, 5 hexagonal faces)
3. *football* (6.3k particles, D3 symmetry, 32 triskelions, 24 C1 HPP vertices, 2 C3 vertices, 6 C1 HHP, 6 hexagonal faces)
4. *D6 barrel* (8.28k particles, D6 symmetry, 36 triskelions, 24 C1 HPP vertices, 12 C1 HHP vertices, 8 hexagonal faces)
5. *tennis ball* (10.2k particles, D2 symmetry, 36 triskelions, 24 C1 HPP vertices, 12 C1 HHP vertices, 8 hexagonal faces)

The C2 basket geometry was delineated from a pool of football particles that were classified into two classes using CISTEM, with the D3 map as the initial model and no symmetry applied. As a last step, to prepare cage maps and metadata (Euler angles) for subparticle analysis, the Euler angles were refined in CISTEM with no symmetry applied, then the maps were reconstructed with symmetry in RELION to enable consistent subtraction of projections from images during the subparticle analysis (refer to the next section) and also to adjust the grayscale of the maps to that of the particle images.

### Subparticle analysis

For each of the five cage geometries, three sets of subparticles were defined: vertices, pentagonal faces, and hexagonal faces. There are 12 pentagonal faces in any of the given cage geometries, but the number of hexagonal faces increases with the diameter of the cage. Vertices are divided to three subcategories. The most abundant vertex geometry in any cage geometry is one made by two pentagonal faces and one hexagonal face (HPP). The other two vertex geometries (PPP, HHP) are either absent in some of the cages or vary in number between different cages.

All the subparticles were extracted using the *localized_reconstruction*.*py* script (*32*) implemented in SCIPION (*48*) and adopted to work with RELION3 rather than RELION1.4. The input parameters for the localized reconstruction script included: 1) STAR file of refined particle images of a cage, 2) a map of the corresponding cage, missing the subparticle of interest, to be projected and subtracted from particle images, 3) C1 symmetry for both particles and subparticles, 4) coordinates of the *center* of the subparticles in *CMM* format, determined such that the clathrin heavy chain part of the subparticle is always centered in its new subparticle box, 5) subparticle box size of 600 pixels (unbinned), 6) Z-axis alignment enabled. The subparticles were extracted from the parent cage particle image. Stacks of both subtracted and non-subtracted subparticles images were created, both unbinned and binned by 2. The subtracted images were only used for high resolution refinement of clathrin at vertices. First, subparticles were re-centered to eliminate centering errors of the extraction step using CRYOSPARC 2D classification and alignment. Then subparticles were refined in CRYOSPARC against an initial model extracted from the corresponding cage map. The only symmetry applied for subparticle refinement was C3 for PPP vertices.

### Heavy chain refinement to high resolution

CRYOSPARC maps and metadata were exported and refined further in CISTEM. To achieve a higher resolution, we focused on the asymmetric vertex of the mini-coat, since the mini-coat cage reconstruction had the highest resolution among cages. At this point the images were unbinned and the super resolution pixels were being used in refinement. After refinement in CISTEM, vertex map and metadata were exported to RELION for CTF refinement where per particle defocus and beam tilt were refined. Afterwards, the final map was generated by refinement in CISTEM.

### Adaptor classification and analysis

After refinement of clathrin heavy chain (and light chain) of hexagonal and pentagonal faces of each cage geometry in CRYOSPARC, the faces were classified individually in RELION into two classes. Centered at β-propellers, a spherical mask with a diameter of 196 Å was applied such that the hypothetical space occupied by adaptors below β-propellers and the inner distal and proximal clathrin legs in that face were all weighted by one, and everything outside the spherical mask weighted by zero with a Gaussian falloff at the edge of the mask. In that manner, 10 hexagonal and pentagonal subparticles were classified to 20 classes initially. 60k hexagonal face particle images from the mini-coat were first classified to two classes. Each class contained 30k particles and both resolved a similar density that resembled the AP2 core density. Therefore, each of the classes were classified further into two classes. Again, the same density was resolved with slightly more features and in a seemingly different orientation. Two more classification steps revealed the AP2 core density presented in Figure 5. A similar hierarchical classification scheme was carried out for the pentagonal face of the mini-coat. Unlike the hexagonal face, no adaptor density was resolved for the pentagonal face.

### Model Building

An initial model for *Bos taurus* CHC and CLC was created from the previously published cryo-EM clathrin model (6SCT (*14*)) from *Sus scrofa*. The sequence for CHC in *Bos taurus* is identical to *Sus scrofa*. The CLC sequences are 96.5% identical, and the nonidentical *Sus scrofa* residues were converted to *Bos taurus* ones by mutating them in UCSF Chimera (*49*). The homology model for a single leg of the triskelion was rigid body fit into our asymmetric vertex map. The density for hydrophobic interactions between the heavy chain and the light chain in conjunction of other bulky side chain densities were in good agreement with the model. Structural differences were noted for CLC residues 188-228 between our map and the porcine map, so these were remodeled in 6 residues steps using Rosetta (*50*) enumerative sampling (RosettaES) and residues 189-199 from 6SCT model as initial model. On each step a larger section of the map was segmented allowing for the 6 residues to be model into the new density. Additionally, compared to the porcine map, we resolved additional CHC tripod residues. Using RosettaES and residues 1596-1626 from 6SCT, 15 additional residues were modeled. For the remaining legs of the triskelion, the leg structure that we modeled as described above was rigid body fit into the corresponding densities. Since our vertex density was not symmetric only the proximal domains of the model fit perfectly into the density. The atoms lying outside of the map were moved into the density using Rosetta torsion space refinement that generally preserves secondary structure and side chain interactions. After the complete vertex model was generated, it was refined using Rosetta cryo-EM refinement protocol resulting in three thousand refined models. The models were sorted by total energy and the model with the lowest energy was selected for further refinement using phenix.real_space_refine (*51*).

Molecular graphics images were produced using the UCSF Chimera package (*49*) from the Computer Graphics Laboratory, University of California, San Francisco (supported by NIH P41 RR-01081).

