## supplemental figures for "The Structures of Natively Assembled Clathrin Coated Vesicles"

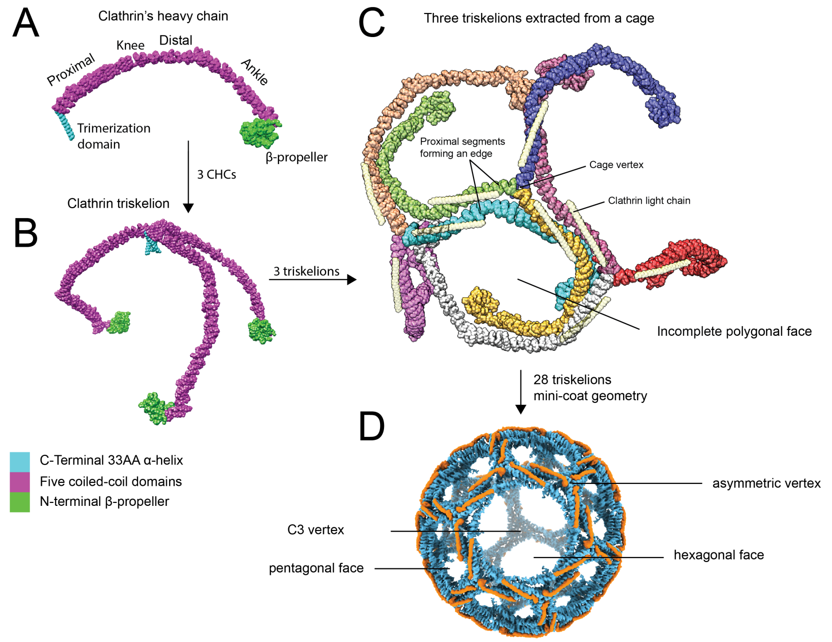


Figure S1. Assembly of clathrin heavy chain at different stages. Color code for panels A and B is the same and different from C and D. A) A single leg-shaped clathrin heavy chain is represented with the names of the domains that each engage in specific interactions. B) Three clathrin heavy chains form a triskelion by interacting through the long α-helical trimerization domain. C) Each CHC has a single color, CLC is represented in light yellow. Interactions between different domains are shown for three triskelions. Two anti-parallel proximal legs make the top layer of an edge, and two anti-parallel distal legs make the bottom layer of an edge. Each triskelion constitutes the center of a vertex, while three knee domains pass near the center, and three distal legs pass under the center of the vertex. Light chain's heptad repeat lies against the proximal domain. β-propellers lie inside polygonal faces. D) Cryo-EM map of the mini-coat. CLC is shown in orange and CHC is shown in blue. Different geometrical elements of the structure are highlighted. There are 4 hexagonal faces, 12 pentagonal faces, 24 asymmetric vertices, and 4 C3 vertices in the mini-coat structure.

A

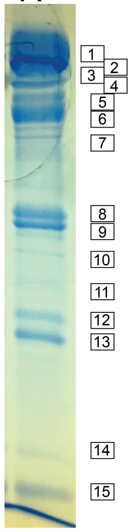


| Gel band | Identified Protein | Accession # | Total Spectra |
|---|---|---|---|
| 1 | Synaptosomal-associated protein, 91kDa homolog | DAA26365.1 | 69 |
| 2 | AP2-associated protein kinase 1 isoform X1 | XP_015320883.1 | 62 |
| 3 | Clathrin heavy chain 1 | CLH1_BOVIN | 482 |
| 4 | Sodium/potassium-transporting ATPase subunit alpha-1 | AT1A1_BOVIN | 49 |
| | AP-3 complex subunit beta-2 | NP_001230264.1 | 20 |
| 5 | AP-2 subunit alpha-1 | NP_001178369.1 | 94 |
| | Amphiphysin isoform X2 | XP_010802730.1 | 23 |
| 6 | AP-1 subunit beta-1 | AP1B1_BOVIN | 118 |
| | AP-2 complex subunit alpha-2 | AP2A2_BOVIN | 105 |
| | AP-1 subunit beta-1 isoform X1 | XP_005218100.1 | 40 |
| | AP-1 subunit gamma-1 isoform X1 | XP_005218546.1 | 17 |
| 7 | Epsin-4 | EPN4_BOVIN | 42 |
| | AP-3 subunit delta-1 | AP3D1_BOVIN | 4 |
| 8 | Tubulin alpha-1D chain | TBA1D_BOVIN | 62 |
| 9 | AP-2 subunit mu | AP2M1_BOVIN | 75 |
| | AP-1 subunit mu-1 | AP1M1_BOVIN | 4 |
| 10 | AP-3 mu3A subunit | AP3M1_BOVIN | 3 |
| 11 | Synaptophysin | DAA12827.1 | 6 |
| | Adaptin ear-binding coat-associated protein 1 | NECP1_BOVIN | 23 |
| 12 | Syntaxin-1B | STX1B_BOVIN | 18 |
| | Syntaxin-1A | NP_001077267.1 | 5 |
| 13 | Clathrin light chain B | CLCB_BOVIN | 88 |
| 14 | Ferritin light chain | FRIL_BOVIN | 74 |
| 15 | Ferritin heavy chain | FRIH_BOVIN | 151 |
| | AP-2 complex subunit sigma | AP2S1_BOVIN | 15 |
| | AP-1 subunit sigma-1A | AP1S1_BOVIN | 7 |
| | AP-1 subunit sigma-1B | AP1S2_BOVIN | 7 |
| | Synaptobrevin-2 | VAMP2_BOVIN | 29 |

B

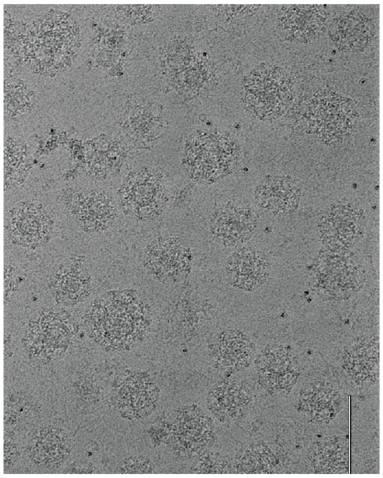

Figure S2. Natively assembled CCVs purified from cow brain. A) On the left, an SDS-PAGE gel lane of the crude sample is shown with bands numbered from high to low molecular weight. In the table, for each gel band, the identity with the highest normalized total spectra count is listed. Mass spectrometry of the crude sample in solution (not shown) agrees with the gel-band analysis. B) An exemplar micrograph of natively assembled CCVs at 33k magnification. Scale bar is 500 nm. Dark spots are ferritin molecules that co-purified in the centrifugation process.

A B C

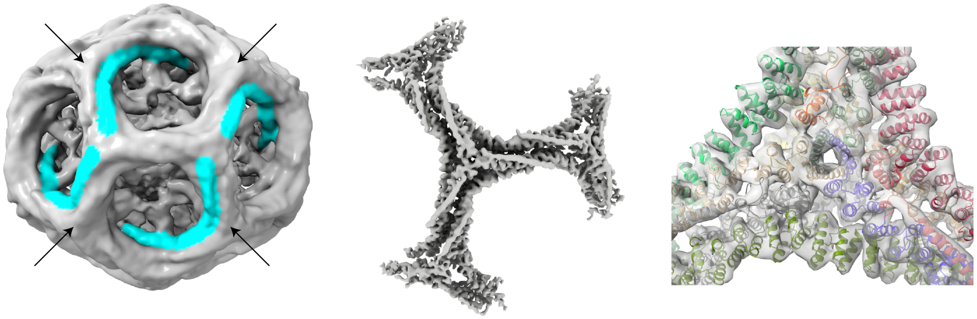

Figure S3. Novel C2 basket geometry with 30 triskelions. A) Arrows show vertices where the head-to-tail exclusion rule is violated. The CHCs highlighted in cyan have a weaker density compared to the rest of the structure. B) Subparticle refinement of the asymmetric HPP vertex of the C2 basket at 9.4 Å. C) Fitting of the generated atomic model for the asymmetric vertex of mini-coat into the asymmetric vertex of the C2 basket (transparent gray) shows no deviation from standard CHC conformations.

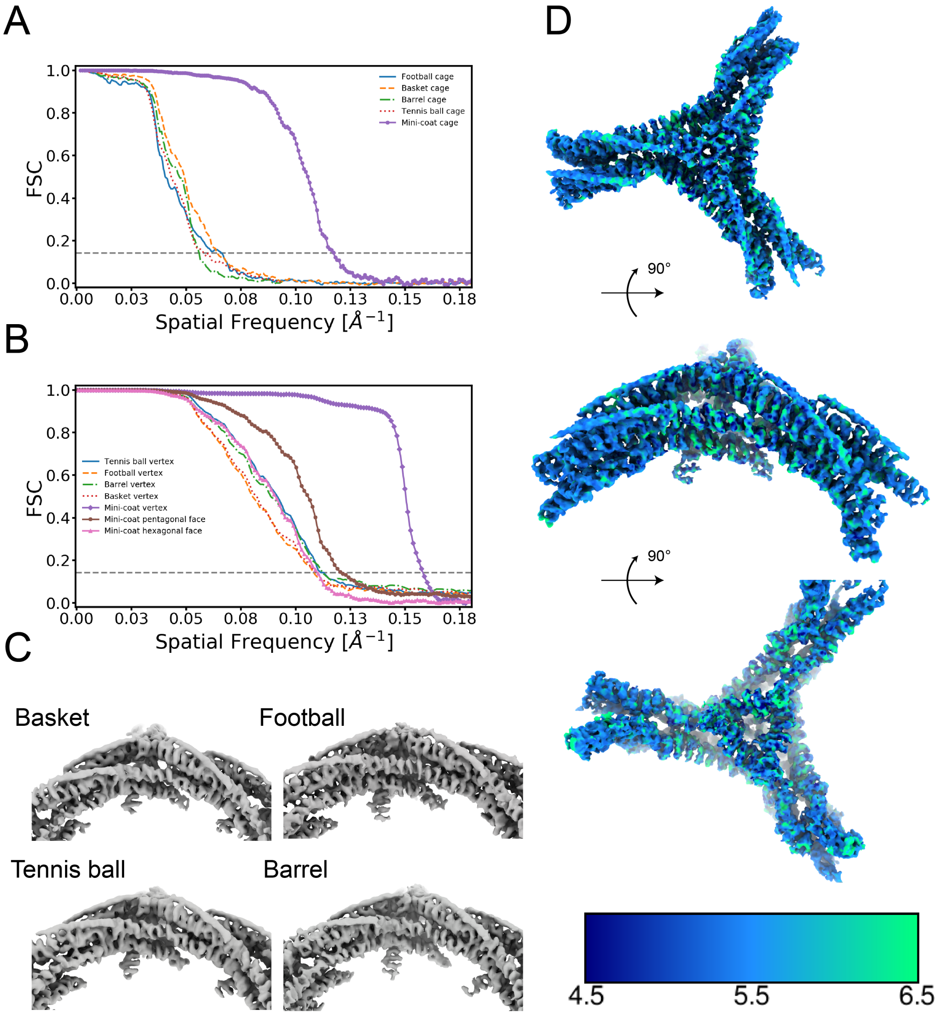


Figure S4. Single particle analysis of cages and subparticles. A) $FSC_{0.143}$ values for cage reconstructions are: football 15.1 Å, basket 15.6 Å, barrel 17.9 Å, tennis ball 17.2 Å, and mini-coat 8.5 Å. B) $FSC_{0.143}$ values for subparticle reconstructions are: tennis ball vertex 8.97 Å, football vertex 9.24 Å, barrel vertex 8.9 Å, basket vertex 9.13 Å, mini-coat vertex 6.3 Å, mini-coat hexagonal face 9.1 Å, mini-coat pentagonal face 8.27 Å. C) asymmetric vertex reconstructions. D) local resolution map for the mini-coat vertex generated by monores. The indicated resolutions are color-coded on the map.

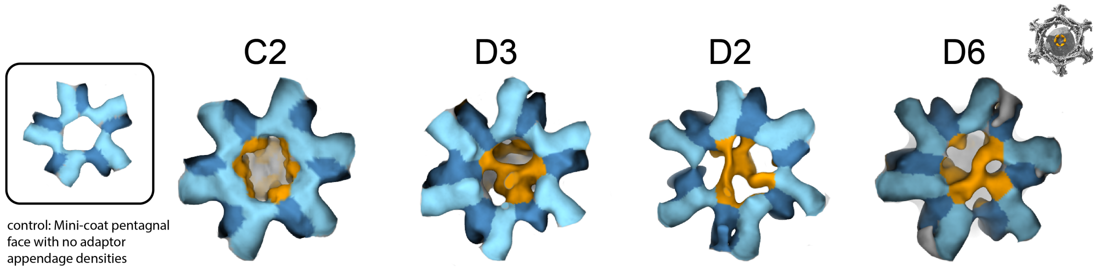


Figure S5. Adaptor appendage densities in hexagonal faces of all cage geometries. At the top right corner, the orientation of all faces is indicated with the hexagonal face of the mini-coat in gray and its β2-appendage in orange. The inset at the left shows a pentagonal face with no adaptor appendages for comparison. The color code is the same as that in Figure 4. β-propellers are in dark blue, the ankle domain is in light blue, and adaptor appendages are in orange. C2 basket, D3 football, D2 tennis ball, and D6 barrel hexagonal faces are shown from left to right, respectively.
